# Distinct striatal subregions and corticostriatal connectivity for effort, action and reward

**DOI:** 10.1101/2020.02.12.925313

**Authors:** Shosuke Suzuki, Victoria M. Lawlor, Jessica A. Cooper, Amanda R. Arulpragasam, Michael T. Treadway

**Author notes:** Correspondence to: Michael T. Treadway, Department of Psychology, Emory University, Atlanta, GA 30306, e.

## Abstract

The ventral striatum is believed to encode the subjective value of cost/benefit options; however, this effect has strikingly been absent during choices that involve physical effort. Prior work in freely-moving animals has revealed opposing striatal signals, with greater response to increasing effort demands and reduced responses to rewards requiring effort. Yet, the relationship between these conflicting signals remains unknown. Using fMRI with a naturalistic, effort-based navigation paradigm, we identified functionally-segregated regions within ventral striatum that separately encoded action, effort, and discounting of rewards by effort. Strikingly, these sub-regions mirrored results from a large-sample connectivity-based parcellation of the striatum. Moreover, individual differences in striatal effort activation and effort discounting signals predicted striatal responses to effort-related choices during an independent fMRI task. Taken together, our results suggest that a dorsomedial region primarily associated with action may instead represent the effort cost of actions, and raises fundamental questions regarding the interpretation of striatal “reward” signals in the context of effort demands.

## Introduction

Weighing the costs and benefits of actions is critical for everyday decisions. The ventral striatum (VS) is widely recognized as a central hub for processing cost/benefit information^1-3^. This role for VS has been supported by robust findings from animal models implicating striatal dopamine (DA) signals in valuation and motivating actions for rewards^4-8^. Building upon the rich preclinical literature, human neuroimaging studies have yielded relatively homogenous results regarding the involvement of the VS in encoding subject-specific, discounted value signals^1^. For example, fMRI studies have demonstrated that VS responses not only are consistently greater for immediate compared to delayed rewards^9-11^, but also explicitly track the delay-discounted subjective value rather than the objective magnitude of rewards^12-14^. Similarly, VS activity is strongly associated with expected reward such that less probable rewards elicit decreased responses^15,16^, and tracks with the probability-discounted subjective value^13^. Moreover, it has been proposed that VS encodes subjective value signals that are domain-general, such that different types of cost/benefits are incorporated and represented on a common scale^17^. Consistent with this, VS has been shown to track subjective value regardless of whether the reward was probabilistic or delayed^13^. Thus, the representation of subjective value signals in the VS detected during cost/benefit decision-making has been largely consistent with the vast preclinical literature highlighting the role of striatal signaling in valuation.

A surprising exception has been the neural representation of costs related to effort (i.e., the amount of work required to obtain rewards). Researchers have used various effort-based decision-making paradigms to decode effort-related value signals^14,18-31^ with widely varying study designs (e.g., timing at which effort/reward information is presented, type and magnitude of effort demands, inclusion of effort performance during the task, inclusion of a learning component). Critically, a majority of these studies did not find a significant association between subjective value and VS activity^14,19-21,23-25,27-30^, not only in humans but also in rodents^32^. This observation is strikingly at odds with prevailing theories about the functional significance of VS, and discordant with consistent patterns of value-related VS activity found in other forms of cost/benefit decision-making^12-14^ as well as evidence demonstrating that disrupting the VS reliably impairs the willingness to work for rewards^7,33,34^.

Recent work examining striatal function in freely-moving animals suggests one potential resolution to this discrepancy^35-38^. Such studies have found that DA inputs to the striatum signal the initiation of vigorous action (“effort activation”)^38^, while also representing the value of cost/benefit choices that are discounted by effort-related costs (“effort discounting”)^6,36^. These observations raise the possibility that detection of effort-discounting signals may perhaps be hindered by the simultaneous presence of an effort-activation signal in most neuroimaging paradigms, and that critical aspects of striatal function may only be revealed by carefully parsing these opposing influences.

Importantly, these action- and value-related signals associated with effort demands may arise from distinct neural populations within the striatum^39,40^. For example, prior work examining the neural substrates of goal-directed behavior has suggested that morphological subregions of the VS, including the nucleus accumbens (NAcc) core and shell^41^, have different functions that together guide actions for rewards^2,42^. Such studies have provided evidence suggesting that neighboring regions within the VS have separable roles in processing reward- and effort-related information^43-46^. In a separate line of work, striatal subregions have also been identified by parcellating the striatum based on intrinsic functional connectivity MRI (fcMRI)^47^, although the functional significance of these subregions remain unclear. Importantly, these reported morphological and connectivity-based striatal organizations raise the hypothesis that effort activation and discounting signals may originate from different subregions within VS; however, this has previously not been tested.

Additionally, the importance of including a naturalistic action component within experimental paradigms is highlighted by evidence suggesting that (i) midbrain DA activity controls the initiation of future action^35^ and (ii) reward-related striatal DA inputs are attenuated in the absence of action initiation^38^. To date, however, most studies investigating the neural mechanisms of effort-based choice in humans have commonly focused on evaluating neural responses during the presentation or outcome of effort- or reward-related cues or choices—but not during periods of effortful action—as the primary method with which to identify the neural substrates of subjective value^14,18-31^. More importantly, these prior studies have not been designed to clearly distinguish effortful or vigorous action from mere movement. This raises fundamental questions regarding the interpretation of striatal “reward” signals given the potentially-critical role of dynamic action. Notably, it has been demonstrated that naturalistic action within virtual-reality paradigms can invoke midbrain DA neuron activity in the mouse brain^48^, offering a means by which this preclinical work can be translated to human neuroimaging.

Here, we combined functional magnetic resonance imaging (fMRI) with virtual navigation to evaluate effort-, action-, and reward-related activity in the human striatum. Specifically, participants navigated through 3D mazes with the objective of obtaining rewards (Fig. 1A), and completion of the mazes varied across four conditions: Two in which individuals actively controlled navigation using (i) high-effort and (ii) low-effort button pressing, and two in which individuals passively completed navigation, one (iii) with motion and the other (iv) without motion. Neural activity during the maze-navigation task was additionally examined in relation to neural responses during a second effort based-benefit decision-making task, which was independently assessed using a paradigm previously published by our group^30^. During this second task, participants made choices about their willingness to perform manual button presses for monetary rewards. As such, we were able to examine neural responses to periods of effort activation and effort discounting, and then see how they contributed to neural responses when making decisions about the value of work.

**Fig. 1.**
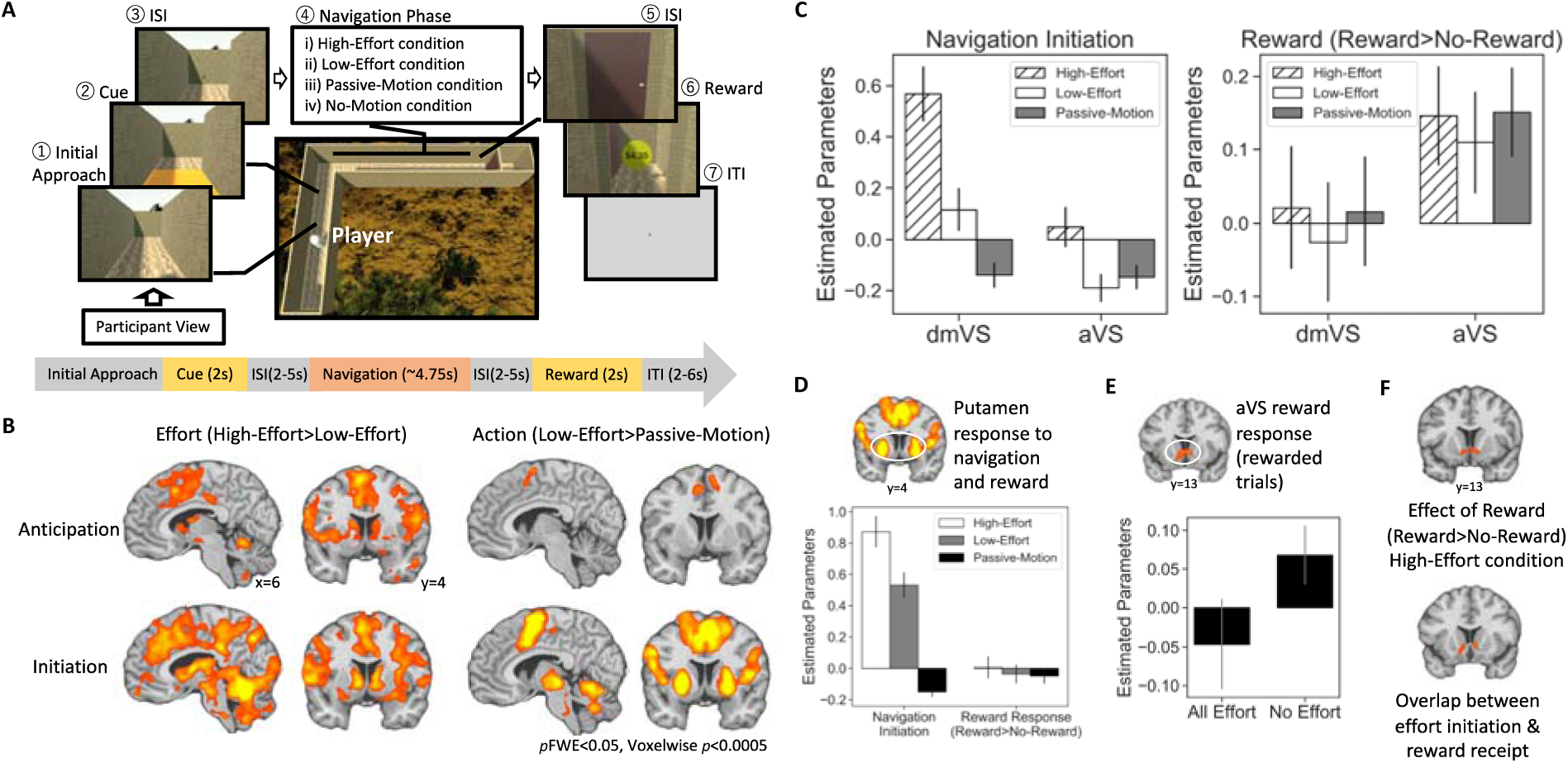
Maze-navigation task schematic and results. (A) Schematic of a task trial. (B) Whole-brain results examining the effect of effort and action during anticipation and initiation of navigation. Maps are *p*<0.05 familywise-error (*p*FWE) cluster-corrected with voxel-threshold *p*<0.0005. (C) Independently-defined ROI analyses showing functional dissociation between the effort- and reward-related VS subregions. (D) Putamen response to navigation and reward. (E) Comparison of reward effects in aVS between effortful (High-/Low-Effort) and non-effortful conditions (Passive-/No-Motion). (F) VS response to reward after effortful navigation and overlap between effort initiation and reward effects.

## Results

### Activity in ventral striatal subregions are modulated by anticipation and initiation of vigorous action

To examine whether the VS encoded *effort activation* in the maze-navigation task, we compared hemodynamic response signals between a High-Effort condition that required rapid and repeated button-pressing to move through the maze, and a Low-Effort condition that allowed individuals to simply hold down buttons to move. This signal was compared at two separate timepoints: (i) when participants learned about the required effort (i.e., anticipation), and (ii) when participants began execution of action (i.e., initiation). Indeed, both the anticipation and initiation of High-relative to Low-Effort increased activity in a dorsal subregion of the VS extending up into dorsomedial striatum and caudate body (MNI coordinates [x,y,z], anticipation: left sub-peak: −12, 0, 15; right sub-peak: 12, 12, 6; initiation: left sub-peak: −9, 0, 6; right sub-peak: 9, 6, −3; Fig. 1B). Notably, this effort-related dorsomedial subregion of VS (“dmVS”) was not modulated by the anticipation or initiation of action alone (i.e., during low-effort navigation), as evaluated by comparing the Low-Effort condition to a Passive-Motion condition that involved automatic progression through the maze without any required finger movement. Instead, action initiation strongly recruited a distinct striatal subregion in bilateral putamen for both the High-and Low-Effort conditions (left sub-peak: −24, 0, −6; right sub-peak: 24, 3, −3). In a follow-up region-of-interest (ROI) analysis using independently-defined ROIs (see Methods), we found evidence for a double-dissociation such that responses within the dmVS (Fig. 1C) and putamen (Fig. 1D) were selective to effort and action, respectively (Region*Effect interaction: *F*_(1,28)_=43.34, *p*<0.001, *η*^2^=0.61).

### Effort discounting is represented in ventral striatal activity during reward receipt

We then examined whether VS also encoded *effort discounting*. Upon completion of mazes, participants were presented with a reward amount ($0.00-$5.00) that varied independently of the navigation condition. An anterior/ventral subregion of VS (“aVS”) responded to reward receipt (left sub-peak: −3, 15, −3, right sub-peak: 6, 15, −3), and was positively correlated with trial-by-trial reward magnitude (left sub-peak: −6, 18, −6; right sub-peak: 6, 15, −3) across all navigation conditions, including a No-Motion condition that involved simply waiting for the approximate duration of the maze. Importantly, while aVS exhibited a reward response within the High-Effort condition, aVS response to reward was significantly lower after effortful navigation compared to navigation that required no effort (*t*_(28)_=−1.88, *p*=0.03; Fig. 1E), suggesting evidence of effort discounting. To further isolate effort discounting in the VS, we examined whether there were areas of VS that responded to reward only during the High-Effort condition. This analysis isolated a region of VS that was at the intersection of aVS and dmVS, suggesting a node of interaction between effort activation and effort discounting (Fig. 1F; peak: 6, 0, −9, *k*=72).

### Functional segregation of effort-, reward-, and action-related striatal subregions recapitulate connectivity-based parcellation of the striatum

Mapping together the above results (Fig. 2A), we observed functional segregation of bilateral striatal subregions associated with effort activation (dmVS), reward (aVS), and action (putamen). Critically, the neighboring regions associated with effort and reward showed clear evidence of double dissociation (Region*Condition interaction: *F*_(1,28)_=13.77, *p*<0.001, *η*^2^=0.33), with dmVS responding strongly to the anticipation and initiation of effort, but not receipt of reward, while aVS responded strongly to the magnitude of reward and appeared to discount reward values by effort, but showed significantly less engagement to anticipation or initiation of effortful action (Fig. 1C; also see Fig. S1 for results using unsmoothed data). Strikingly, this functional segregation appeared to largely overlap with a previously-reported connectivity-based parcellation of the striatum (Fig. 2A)^47^. To validate the apparent functional distinction between these two VS subregions, we compared patterns of functional connectivity during resting-state using each of the functionally-defined and connectivity-based striatal subregions as seed regions. Indeed, despite only partial overlap of the ROI definitions derived from our task and the prior parcellation study^47^ (*Sørensen*–*Dice*: aVS: 0.19; putamen: 0.40; dmVS: 0.20), we observed highly overlapping connectivity profiles (Fig. 2B), suggesting that these ventral striatal subregions participate in distinct functional networks. These connectivity profiles were additionally replicated in an independent resting-state dataset collected with high-resolution (7-Tesla) fMRI (Fig. 2B)^49^.

**Fig. 2.**
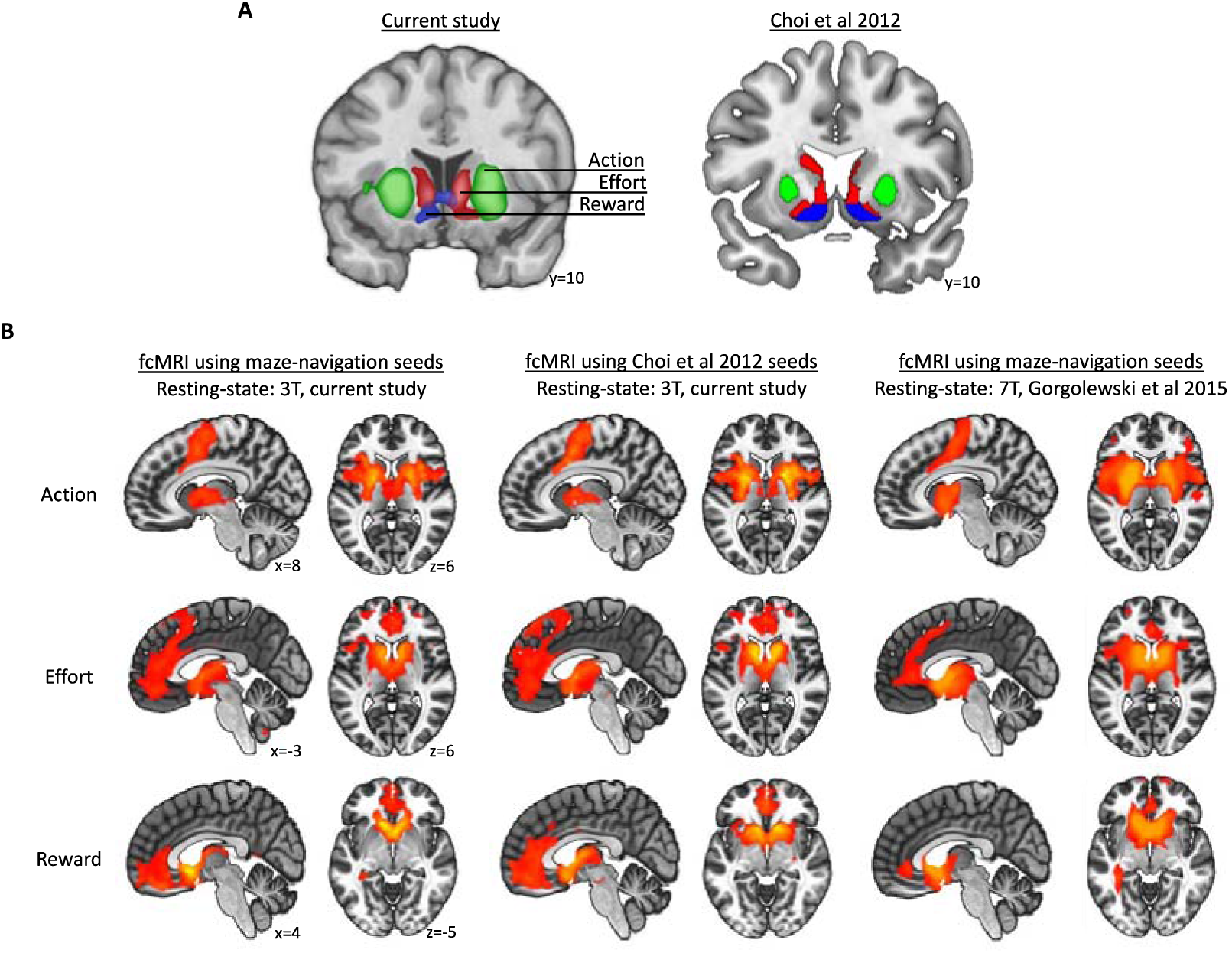
Functional segregation of striatum and comparison of connectivity profiles with previously-identified striatal parcellation based on intrinsic connectivity. (A) Striatal subregions identified in the current study and corresponding subregions identified via connectivity-based parcellation^47^. (B) Comparison of seed-to-voxel functional connectivity maps using striatal regions in (A) as seed regions and resting-state data from the current study and prior work^49^. Maps are *p*<0.05 familywise-error corrected with voxelwise *p*<0.0005.

### Lack of ventral striatal activity related to value during effort-based decision-making paradigm

To test the hypothesis that effort activation and discounting signals may interfere with the detection of subjective signals during effort-based decision-making, we also measured neural activity during a second fMRI paradigm in which participants made a series of binary choices based on the presented amount of reward and effort required for each option (Fig. 3A). Importantly, the effort and reward amounts were presented sequentially in attempt to isolate an effort-activation signal during the anticipation of various effort demands. Replicating our previous findings using this same task^30^ as well as results from other fMRI effort-based paradigms, we did not find an association between VS activity and subjective value of the chosen option (Fig. 3B; Table S2). Even when using multivoxel pattern analysis, voxels in the VS were unable to classify whether presented information reflected reward or effort information (Fig. S2). In contrast, classifiers trained on activity across all brain voxels successfully decoded whether reward or effort information was presented for 78.9% of individuals (Fig. S3). As described above, the absence of such an effect is surprising, given the robust activation of VS by subjective values derived from cost/benefit decisions involving other categories of response costs (e.g., delay, risk, or loss).

**Fig. 3.**
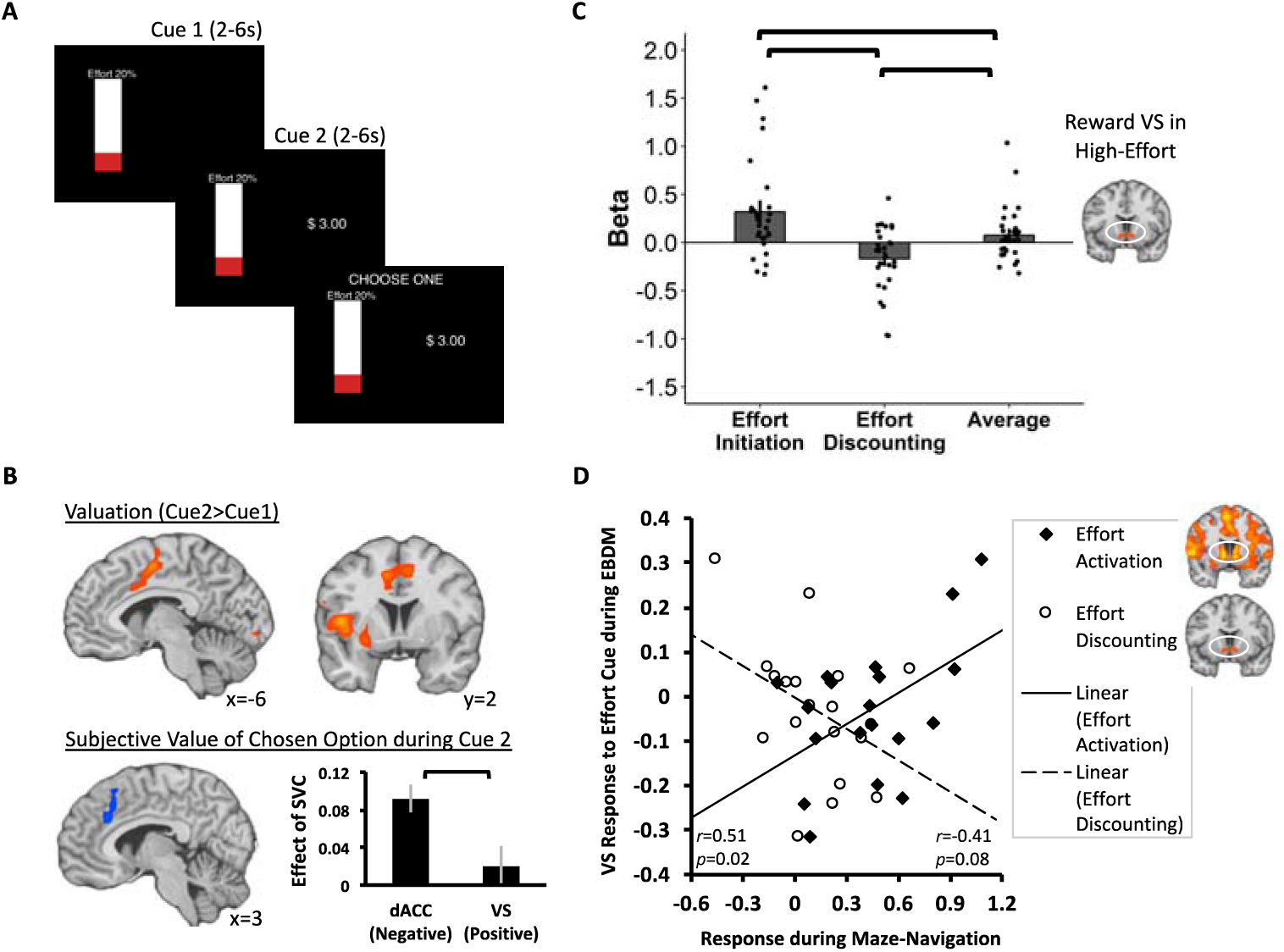
Schematic and results of effort-based decision-making task and cross-paradigm analyses. (A) Schematic of a task trial. (B) Whole-brain results replicate our prior findings using this paradigm ^30^, including activation of dorsal anterior cingulate cortex (dACC) and insula at Cue 2, association between dACC and negative subjective value, and absence of VS response across contrasts. Maps are *p*<0.05 familywise-error corrected with voxelwise *p*<0.0005. (C) aVS encodes both effort activation (High-Effort>Low-Effort at navigation initiation) and effort discounting (High-Effort>Low-Effort at reward receipt) that average to a null-group effect. (D) Effort-related striatal signals during maze-navigation predict VS responses to effort information during effort-based decision-making (EBDM). Error bars in bar plots represent standard error of the mean.

### Ventral striatal activity is modulated by opposing effects of effort initiation and effort discounting

One possible explanation for the absence of effort-related subjective-value encoding in VS is that when response costs involve physical effort, opposing effort-activation and effort-discounting signals within VS may impede the ability to detect overall VS responses. To test this possibility in the maze-navigation task, we tracked responses within the VS region that responded to reward after individuals completed effortful navigation (Fig. 1F). This revealed two distinct patterns of effort-related activity in the VS depending on the timepoint, as evidenced by a significant Condition (High-Effort, Low-Effort)*Phase (navigation initiation, reward receipt) interaction (*F*_(1,28)_=16.72, *p*=0.0003, *η*^2^=0.37): VS responses were (i) significantly greater when individuals began executing effortful action compared to mere action (*t*_(28)_=3.42, *p*=0.002), reflecting an effort-activation signal, and (ii) significantly lower during reward receipt after individuals have expended effort compared to less effort (*t*_(28)_=−2.66, *p*=0.01), reflecting an effort-discounting signal. Importantly, averaging the opposing effects associated with effort activation and effort discounting within individuals led to a null group-level effect (*p*=0.16; Fig. 3C).

However, such “averaging” is clearly artificial, as these task conditions did not occur simultaneously. A much stronger test would therefore be to show a mix of effort activation and effort discounting signals in response to the same effort-related stimulus. We therefore extracted the “effort activation” and “effort discounting” signals from the maze-navigation task, and used them as predictors of neural responses to effort cues in our second (independent) task. Interestingly, individual differences in effort-activation signal were positively associated with VS responses during presentation of effort information during decision-making (*r*_(17)_=0.51, *p*=0.02), whereas individual differences in effort-discounting signals exhibited the opposite pattern (*r*_(17)_=−0.41, *p*=0.08; Steiger’s *Z*=2.88, *p*=0.004; Fig. 3D). These findings provide a potential explanation for the longstanding discrepancy regarding the involvement of VS in encoding subjective value during effort-based decision-making epochs, as such epochs might reasonably be expected to elicit both effort-discounting and effort-activation signals.

## Discussion

For almost two decades, neural responses in the ventral striatum have been commonly observed in response to rewards and discounted value of rewards. Thus, it has been surprising that most functional imaging studies in humans and animals have shown weak or inconsistent activation of VS during effort-based choice^14,18-31^. One methodological gap in prior studies is the failure to take into consideration striatal signals during effortful action that have been detected under free-movement conditions in rodents^35,38,50,51^, which may impact the interpretation of reward-related striatal BOLD activity. Such studies have highlighted the presence of striatal signals that are heavily linked to dynamic movement^35,38^, which can also be evoked by movement within virtual-reality paradigms^48^. Thus, the present study used a virtual navigation paradigm and revealed the presence of opposite signals in the VS related to (i) effort activation during the anticipation and initiation of effortful action and (ii) effort discounting during reward receipt after individuals expended effort. Importantly, the current study finds that effortful action is needed to isolate these effects in the VS, as our second fMRI paradigm that relied solely on effort- and reward-*cues* failed to differentiate them, even when using more sensitive multivariate techniques. This observation is critical as prior studies of effort-based decision-making have largely focused on encoding of values at the point of choice or in response to feedback^14,19-21,23-25,27-30^. Critically, by using individual differences in estimates of effort-activation/effort-discounting signals derived from the maze navigation task, we were able to observe their simultaneous, conflicting influences when processing effort-related cues, providing a viable explanation to the general failure to detect clear subjective-value signals in the VS during effort-based decision-making in these prior work. Specifically, our results suggest that increased effort requirements may invoke a greater effort-activation response that may obscure detection of the value-related effort-discounting effect in the VS.

Additionally, the current results provide insight into the functional topography of signals associated with effortful movement and reward. Despite the rich body of research implicating midbrain DA in representing value-related information and invigorating movement^4^, little work has been done to simultaneously examine the striatal representation of these two functions in humans. Here, we identified an aVS region associated with reward, a neighboring dmVS region associated with effort-initiation, and bilateral putamen associated with initiation of simple, low-effort action. Interestingly, we observed an invigoration effect in dmVS during the anticipation and initiation of high- but not low-effort, suggesting that mere action (i.e., making a single choice response) may not be sufficient to shape VS signaling—further highlighting the importance of studying the neural substrates of effort-based decision-making that include effortful action components.

To our knowledge, the VS subregions identified here have not been previously characterized, though their existence is supported by prior large-scale connectivity results^47^. Indeed, we found overlap in functional-connectivity profiles between the striatal subregions identified here and the connectivity-based striatal parcellation^47^. Critically, this finding helps uncover potential functions of these network-based subregions of the striatum. Our results also show similarity to the pattern of results of a recent meta-analysis of subjective-value encoding^1^, where positive subjective value across studies was found to scale with activity in aVS, whereas negative subjective value (i.e., loss or punishment) was associated with activity in an area similar to dmVS. As activity in dmVS was only apparent as a function of effort but not action, whereas motor areas of putament appeared to respond to both increased effort and mere action, it raises the intriguing possibility that dmVS is more specifically involved with encoding negative value or cost associated with effortful actions. Finally, it is worth noting that similar functional heterogeneity within the ventral striatum has been observed in animal models [floresco]. Electrical recordings in non-human primates have found that distinct neuronal populations within the striatum may be responsible for coding action and outcomes^39,40^. In rodents, while both the NAcc core and shell are involved in Pavlovian learning of stimulus and outcome (e.g., cue and reward), they play markedly different roles in terms of mediating Pavlovian influences on instrumental behavior^42,52^.

Taken together, our results suggest that dmVS activity at the initiation of effortful action is distinctly associated with neural representations of effortful action relative to reward-responsive aVS. Importantly, our results suggest that simultaneous and conflicting influences of invigoration and value in these neighboring regions may confound the absence of subjective-value signals in prior work on effort-based decision-making. It is further notable that these distinct sub-regions of ventral striatum were only observable when using a naturalistic paradigm, echoing recent discoveries of striatal DA function found in freely-moving animals^35-38^. In sum, these data will help advance our understanding of how the ventral striatum distinctly encodes effort-related costs, and highlights the value of naturalistic and dynamic paradigms for achieving a deeper understanding of real-world brain function^53^.

## ACKNOWLEDGEMENTS

This work was supported by funding from the NIMH R00 MH102355 and R01 MH108605 to MTT and F32 MH115692 to JAC, and the National Science Foundation Graduate Research Fellowship Program DGE-1444932 to ARA. The authors would like to thank Johsua Buckholtz and Daniel Dilks for helpful discussion and commentary. The authors additionally thank Brittany DeVries, Makiah Nuutinen, Emma Hahn, Danielle Harrison, Annabel Lu, Maryam Rehman, Jeffrey Yang, Kristi Kwok, Samuel Han, and Nimra Ahad for their assistance in data collection.

## AUTHOR CONTRIBUTIONS

SS wrote the first draft of the paper; SS and MTT designed the research; SS performed the research; SS, VL, JAC, ARA, and MTT analyzed the data; SS, VL, JAC, ARA, and MTT wrote the paper.

## Methods

All procedures were approved by the Emory Institutional Review Board. We recruited 30 healthy adults from the Atlanta community through fliers and online advertisements. Eligibility was determined using an online pre-screening survey. Participants were eligible if they were right-handed, English-speaking individuals between the ages of 18 and 35. Exclusion criteria included: (1) MRI contraindications (e.g., claustrophobia, metallic implants, CNS diseases, pregnancy in females); (2) Current use of psychoactive medications, investigational drugs, or those that affect blood flow (e.g., for hypertension); and (3) Current medical, neurological, or psychiatric illnesses.

Participants first provided informed consent and completed the MRI safety form. Prior to MRI scanning, participants underwent a training and calibration procedure on the maze-navigation task, and on the effort-based decision-making task for a subset of the participants (n=19). In the MRI scanner, all participants completed the maze-navigation task (n=30), and two subsets of the participants completed the effort-based decision-making task (n=19) and/or a resting state scan (n=19). After scanning, participants were taken to a separate interview room to complete a debriefing interview. Participants were compensated for their participation upon completion of study procedures.

Imaging data acquired from one participant during the maze-navigation task was excluded prior to analysis due to the individual falling asleep during the task. Thus, the final sample included 29 participants for the maze-navigation task analysis (M_age_=24.41; SD_age_=5.43; 21 female), 19 participants for the effort-based decision-making task (M_age_=24.26; SD_age_=4.64; 14 female), and 19 participants for the resting-state scan (M_age_=24.94; SD_age_=5.53; 12 female).

### Maze-navigation task

The task was programmed using Unity 3D (Unity Technologies ApS). A schematic of a single trial is shown in Figure 1A. On each trial, participants completed first-person navigation through a single-path virtual maze in pursuit of monetary rewards. Each trial was associated with one of four maze structures, two of which required a single 90° turn (left or right), two of which required two 90° turns (left-then-right or right-then-left), and all of which were equated in approximate navigation time (∼5s). Regardless of structure, each maze was comprised of 1×1 unit^2^ floors placed adjacently to form a path, bounded by 1×1 unit^2^ walls. Participants controlled movement using a 4-button box with the right (i.e., dominant) hand. Specifically, the middle finger button moved the participant forward (r=2.2units/s), and the index and ring finger buttons each rotated the player clockwise and counterclockwise, respectively (ω=0.5π rad/s). Holding down the buttons resulted in continuous movement or rotation (except during the High-Effort condition, as detailed below). Acceleration was applied to player motion to mimic real-life motion. Only one type of motion was allowed at any given moment (i.e., pressing multiple buttons resulted in no motion). The participant view was set at 0.6 units above the floor, with a −10° tilt along the axis parallel to the ground.

Importantly, each trial was associated with one of four navigation conditions: i) The *High-Effort* condition required participants to repeatedly press the button associated with forward-motion to advance through the maze. In this condition, each button press moved the participant forward by an incremental distance, and the distance traveled per button press (i.e., effort level) was individually calibrated during the training procedure; ii) The *Low-Effort* condition required participants to advance through the maze with the default button controls (i.e., simply hold down buttons to move and rotate). This condition served as an active-navigation control for the high-effort condition, which allows for examination of effects specifically related to vigor by subtracting out activity associated with mere action; iii) The *Passive-Motion* condition required participants to view moving through the maze without making any action. This condition served as a no-action control for the low-effort condition to examine effects related to mere action; and iv) The *No-Motion* condition required participants to wait for the approximate duration of the maze (∼4.75s), after which the participant was teleported to the goal. This condition served as a visual motion control for the Passive-Motion condition. Thus, two of the conditions (High-Effort/Low-Effort) required individuals to actively navigate through the maze, while the other two conditions (Passive-Motion/No-Motion) entailed passive completion of navigation. Participants failed the trial if they did not reach the goal within a liberal time-limit (5.5s) in the active navigation conditions, or if they initiated action in the passive navigation conditions.

Each trial proceeded as follows: (1) Initial Approach phase: At the beginning of each trial, the participant position was initialized to the beginning of a maze, immediately after which participants could initiate movement. (2) Cue phase: Upon reaching a ‘trigger’ location in the maze, participants were rendered immobile and the floor unit immediately in view changed color for 2s, before returning to its original texture. The colored cue informed participants to the navigation condition for that trial. (3) Jittered interstimulus interval (ISI): The cue was immediately followed by a jittered fixation period, whereby a ‘+’ was rendered on top of the maze scene for a Poisson-distributed jitter period (2-5s) with a mean duration of 2.5s. (4) Navigation phase: After the fixation cross disappeared, participants regained control of movement and completed navigation according to the navigation condition, as detailed above. On failed trials, participants were presented with a feedback screen for 2s indicating that they failed, followed by an intertrial interval (ITI; skip to (8)). (5) Jittered ISI: Upon successfully reaching the door at the end of the maze (the ‘goal’), participants were again rendered immobile for another jitter period (2-5s) with a mean duration of 2.5s. (6) Reward phase: Following the ISI, participants were presented with an animation of the door opening followed by a monetary reward, represented by a dollar amount rendered on the surface of a coin. Each trial was associated with one of 4 bins of reward magnitudes ($0, $1.68-2.78, $2.79-$3.89, $3.90-5.00), from which an amount was randomly selected. (7) Rating phase: Once every 4 trials, starting on the first trial of each run, the participant was asked to make a mood rating on a Likert-scale between 1 (not happy at all) to 4 (very happy) using button press. (8) Jittered ITI: After reward receipt, failure feedback, or mood rating, participants were presented with a ‘+’ rendered on a grey screen for a duration (2-6s) jittered around 3s.

Prior to scanning, participants completed a 15min training procedure of the maze-navigation task. Participants were introduced to the movement controls, navigation conditions, and trial structure, and informed that a proportion of the reward they obtain on each trial will be added to their compensation as a bonus. Once participants indicated understanding of task instructions, they completed 16 practice trials (4 per navigation condition) on a laptop computer. Participants were instructed to use the same hand and fingers that would be used to perform the task in the MRI scanner.

Each participant’s average rate of key pressing during the high-effort trials were used to calibrate their effort levels for the in-scanner task. To ensure collection of button pressing rates for sufficient durations of time, the distance traveled per button press in the high-effort condition during the practice was set low (0.11 units/press) and no time-limits were imposed. However, participants were informed that there would be a time-limit for the active navigation conditions when they complete the task in the scanner and were instructed to complete the mazes during the practice as quickly as possible. From the practice data, the calibrated effort level (i.e., distance, d, per button press) for each participant was calculated as

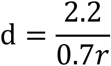

where r is the average key-pressing rate during the first 5.5s of the navigation phase (=time-limit on in-scanner trials) in the High-Effort trials. 2.2 corresponds to the default rate of motion (unit/s). 70% of the average rate was chosen to minimize motion in the scanner, whilst maintaining similar completion times across conditions. All participants successfully completed the practice trials and demonstrated understanding of the four navigation conditions.

During the in-scanner task, the 4 navigation conditions and 4 reward bins were equally distributed and balanced across trials and runs. Each participant completed 3 runs with 32 trials each (∼11min/run), and trials were presented in a fixed-randomized order. Participants successfully completed the maze on 93±2% of trials, indicating low rates of failure. Importantly, mean completion times across conditions were all within 100ms (M_High-Effort_=4.6s; M_Low-Effort_=4.5s, M_Passive-Motion_=4.7s; M_No-Motion_=4.7s), suggesting that condition-based differences in “time-on-task” were unlikely to significantly affect results. Individuals initiated navigation faster on High-Effort trials (M=550ms, std=200ms) compared to Low-Effort trials (M=630ms, std=210ms; *t*_(28)_=2.55, *p*=0.02). Self-reported mood ratings did not differ between the active (High-/Low-Effort) and passive (Passive-/No-Motion) navigation conditions (*p*=0.29; M=2.9, std=0.7).

### Effort-based decision-making task

The task was previously programmed using Psychtoolbox for MATLAB ^30^. A schematic of a single trial is shown in Figure 3A. In this task, participants made a series of binary choices provided information about the dollar amount of the reward and the required amount of physical effort to obtain that reward for each choice option. Specifically, participants chose whether to receive $1.00 for no work or to complete an effortful task for a larger reward of varying amounts. On each trial, the effortful task option was associated with one of 4 bins of reward magnitude ($1.25-2.39, $2.40-3.49, $3.50-4.60, and $4.61-5.73), from which a dollar amount was randomly selected. The effort level was presented as the height of a vertical bar (20%, 50%, 80%, or 100% of the participant’s individually-calibrated maximum effort level).

Prior to the scan, participants completed a training and calibration procedure. Calibration was conducted via three independent trials in which participants were asked to repeatedly press a key using their left (i.e., non-dominant) pinky finger as rapidly as possible within 20s. Each participant’s maximum effort level was operationalized as the average number of keys pressed across the three trials. Participants then practiced completing the effortful tasks that varied in the required amount of key press (20, 50, 80, and 100% of their maximum effort level) within a constant time-limit (20s). They completed 4 trials at each level for a total of 16 trials. Participants were informed that the physical effort component would not be completed inside the MRI scanner, but would be completed immediately following the scan, based on the in-scanner choices. Additionally, participants were instructed that three trials would be randomly selected at the end of the session, from which the reward they earned would be added as a bonus to their compensation.

Each trial was structured as follows: Participants were first presented with either the reward or effort magnitude of the effortful option (Cue 1) on the left or right half of the screen for a jittered duration (2-6s; mean=2.98s). This information remained on the screen while the other piece of information (Cue 2) appeared on the opposite side of the screen for another jittered duration (2-6s; mean=3.23s). Participants were then prompted to choose whether to accept the presented effortful task option, or reject the option in favor of receiving $1.00 for no work (Choice phase). Participants indicated their choice using button press (index-finger button to accept; middle-finger button to reject), which was then presented on the screen with the words “ACCEPTED” or “REJECTED” according to their choice. Order of information (effort-then-reward vs. reward-then-effort) and the presented side of screen were counterbalanced across trials. Each participant completed 2 runs with 44 trials each (∼9min/run), and trials were presented in a fixed-randomized order.

After the scan, participants were presented with each of the effortful choice options they accepted during the in-scanner task and performed the associated tasks. For each trial, participants were given the option to change their choice, to examine whether performing the effortful task in real time influenced their willingness to expend effort. We observed that choice behavior was consistent across contexts, such that participants made the same choice on 95±3% of trials.

### Behavioral and imaging data acquisition

Stimuli were presented via back-projection mirror, and participants completed the maze-navigation and effort-choice task using an MR-compatible 4-button box (Current Designs Inc). Inflatable pads (Multipad 01; Pearltec AG) placed around participants’ heads were used to minimize head motion.

Participants were scanned in a 3-Tesla Siemens TIM Trio scanner (Siemens AG) with a 32-channel head-coil using multiband structural and functional imaging^54^. Each session began with a 3-plane localizer scan for slice alignment, and a single-shot, high-resolution structural MPRAGE sequence (TR/TE=1900/2.27ms; flip angle=9°; FoV=250×250mm; 192×1.0mm slices). Blood oxygen level dependent functional images were acquired with T2*-weighted EPI sequences with a multiband acceleration factor of 4 (TR/TE=1000/30.0ms; flip angle=65°; FoV=220×220mm; 52×3.00mm slices).

### Behavioral data analysis

Behavioral data from the maze-navigation task were used to examine latencies to initiate action, rates of navigation completion, durations of navigation completion, and mood ratings. Latency to initiate action was operationalized as the time between the offset of the jittered ISI following the Cue phase and movement onset, and was computed separately for each of the active navigation conditions (High-/Low-Effort). The rate of navigation completion was calculated as the number of successfully-completed trials divided by the total number of trials. Navigation time was calculated separately for each navigation condition, operationalized as the time between the onset of movement and reaching the goal for the active navigation trials (High-/Low-Effort), and between the offset of the jittered ISI following the Cue phase and reaching the goal for the passive navigation trials (Passive-/No-Motion). Self-reported mood ratings were averaged across trials separately across the active navigation conditions (High-/Low-Effort) and across the passive navigation conditions (Passive-/No-Motion).

Behavioral data from the effort-based decision-making task were used to compute the estimated subjective costs that individuals associate with varying levels of effort. To do this, participants’ choice data were fit to a two-parameter power function used in prior work ^27,30^:

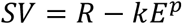

which describes subjective value (SV) as the objective reward magnitude (R) discounted by the subjective cost determined by the effort level (E; between 0-100%) and two free parameters, k and p.

### Imaging data analysis

Functional images from the maze-navigation and effort-based decision-making tasks were preprocessed using SPM12 scripts through NeuroElf v1.1 (neuroelf.net). Specifically, images were co-registered to the structural image, motion-corrected, warped to the MNI template, and smoothed using a Gaussian filter (6mm full width-half maximum). Raw and preprocessed data were subjected to multiple tests for quality assurance and inspected for spiking and motion. Volumes were discarded if the root mean square of motion parameters exceeded a single voxel dimension (3mm), or if striping was identified through visual inspection of each functional volume. Subject-level modeling of trial events was conducted using robust regression to reduce the influence of strong outliers. Effects in group-level whole-brain analyses were considered significant at a voxelwise threshold of *p*<0.0005 combined with a cluster-extent threshold estimated for each contrast map, resulting in FWE-correction of *p*<0.05, unless otherwise specified.

For the maze-navigation task, the subject-level general linear model (GLM) included the Cue phase, first half of the Navigation phase (Navigation-Start phase), and Reward phase as regressors of interest, with each regressor separated by navigation condition. To examine the effect of reward receipt, the Reward phase was further divided into rewarded (reward > $0.00) and non-rewarded (reward = $0.00) trials. A second GLM included the trial-by-trial reward magnitude as a parametric modulator for the Reward phase, instead of separating the rewarded and non-rewarded trials. These models also included the Initial Approach phase, latter half of the Navigation phase, and Rating phase, as well as the ISIs, to omit their influences on the implicit baseline. In addition, motion parameters and their squares, as well as high-pass filter parameters were included as additional nuisance regressors. Given the potential impact of smoothing on the functional segregation of striatal subregions detected in the above models, two other GLMs were generated using unsmoothed images but otherwise following the same procedures as the first and second GLMs.

Group-level contrasts were generated to examine the effects of effort (High-Effort>Low-Effort) and Action (Low-Effort>Passive-Motion) during the Cue and Navigation Start phases, and to examine the effect of reward (Reward>No-Reward, across all conditions) and magnitude of reward (in the parametric models) during the Reward phase. To examine the effect of effort on striatal response to reward, an ROI analysis was conducted by tracking responses within the aVS region found to respond to reward magnitude (left sub-peak: −6, 18, −6; right sub-peak: 6, 15, −3; *k*=77) in the parametric model. Then, we extracted beta parameters during reward receipt on rewarded trials in this independently-defined ROI, and a pairwise comparison was conducted between the conditions that required effort (High-/Low-Effort) and those that did not require effort (Passive-/No-Motion).

To examine the functional selectivity of striatal subregions found during the maze-navigation task, ROI analyses were conducted using a leave-one-subject-out (LOSO) approach to avoid any circularity or non-independence in the definition of ROIs ^55^. This was conducted by defining ROIs using group-level contrasts that include data from all but one participant; extracting beta parameters from the removed participant’s data; and repeating this procedure for all participants. Beta parameters were extracted from striatal subregions associated with effort-initiation (Navigation-Start phase, High-Effort>Low-Effort), action initiation (Navigation-Start phase, Low-Effort>Passive-Motion), and reward receipt (Reward phase, reward>no-reward, across all conditions). Effects of effort, action, and reward for each subregion was calculated by subtracting beta parameters according to the relevant contrasts (Effort: High-Effort – Low-Effort; Action: Low-Effort – Passive-Motion; Reward: Reward – No-Reward, averaged across navigation conditions). A 2 Region (dmVS, putamen) * 2 Effect (Effort, Action) repeated-measures ANOVA was conducted to test for functional dissociation between dmVS and VS, and to examine whether dmVS activity was selective to effort and not mere action. Additionally, a 2 Region (dmVS, VS) * 2 Effect (Effort, Reward) repeated-measures ANOVA was conducted to test for functional dissociation between these neighboring regions associated with effort and reward.

Additionally, to examine the presence of both effort-activation and effort-discounting effects within a VS region, we functionally-defined a VS ROI (peak: 6, 0, −9; *k*=72) that responded to reward after completing effortful navigation (Reward>No-Reward, High-Effort condition). This reward-sensitive ROI specific to the High-Effort condition was defined at a more lenient voxelwise *p*<0.005 given that this region was necessarily defined on only 25% of the data. We note that this threshold was used for ROI definition only and not for subsequent inferential analyses. Beta parameters from this independently-defined ROI were extracted from the High-Effort and Low-Effort conditions during the Navigation Start and Reward phases, and a 2 Condition (High-Effort, Low-Effort) * 2 Phase (Navigation Start, Reward) repeated-measures ANOVA was conducted. Pairwise comparisons were conducted to parse out the nature of significant effects.

The CONN toolbox was used for resting-state fMRI data analysis. Preprocessing of images included motion correction, co-registration to structural scan, MNI normalization, and smoothing using an 8mm Gaussian smoothing kernel. Striatal subregions were functionally-defined from the maze-navigation data and used as seed regions to create individual seed-to-voxel fcMRI maps. Specifically, the seed regions included a bilateral dmVS region associated with effort initiation (Navigation Start phase, High-Effort>Low-Effort), a region in bilateral putamen associated with action (Navigation Start phase, Low-Effort>Passive-Motion), and a bilateral VS region associated with reward (Reward phase, parametric effect of reward magnitude, across all conditions). For each of the three fcMRI maps, comparison maps were generated using the spatially-corresponding subregions among the previously-reported connectivity-based parcellation of the striatum (regions 4, 5, and 7 in the seven subregion parcellation)^47^. Additionally, fcMRI maps using striatal regions defined from the maze-navigation task were computed using an independent set of 7T resting-state fMRI data (N=22) ^49^ downloaded from OpenNeuro (openneuro.org). Specifically, two runs per subject of whole-brain resting-state data from session one in the dataset were included for the purpose of our study. Processing of these data followed the same procedures through CONN as described above, except for using a 6mm smoothing kernel to account for smaller voxel size. Further, *Sørensen*– *Dice* indices were calculated for each ROI defined in the maze-navigation task and the corresponding connectivity-based ROI ^47^ to test for spatial similarity between the ROI definitions. To do this, we used the nilearn—http://nilearn.github.io—library to resample the image space and affine of task-based ROI masks on the connectivity-based ROI masks.

For the effort-based decision-making task, the first subjective-level GLM included the Cue 1 and Cue 2 phases as regressors of interests, with Cue 1 regressors separated by type of information presented (effort vs. reward). We also included the Choice and Rating phases to omit their influence on the implicit baseline. Again, motion parameters and their squares, as well as high-pass filter parameters were included as additional nuisance regressors. At the group-level, we examined regions in which activity was higher during Cue 2 compared to Cue 1 (Cue 2>Cue 1). To examine the effect of subjective value during the valuation period, we additionally included the trial-by-trial SV of the chosen option as a parametric regressor for Cue 2 in a second GLM. At the group-level, we focused on the negative parametric effects of SV to identify regions that increased in response to decreasing value (e.g., higher effort costs). ROI analyses were conducted to examine the effect of negative SV in the dACC and VS. To do this, the dACC ROI (left sub-peak: −10, 26, 32; right sub-peak: 10, 24, 36; *k*=334) was functionally-defined using the same parametric analysis (effect of negative SV) from our prior study using this task ^27,30^ at *p*<0.05 familywise-error corrected with voxelwise *p*<0.0005, and the VS ROI was defined using an anatomical nucleus accumbens mask (left peak: −10, 10, −8, k=309; right peak: 10, 10, −8, k=310). A pairwise comparison was conducted on the extracted parametric effects of negative SV within the dACC and VS.

Multivariate pattern analysis (MVPA) of the effort-based decision making task was performed using the scikit-learn^56^ and nilearn libraries. We trained linear support vector machines with the C parameter fixed at a value of 1. Each classifier used unsmoothed functional images as features to predict the type of information (effort or reward) shown at Cue 1. For each trial, we selected four fMRI volumes offset by four seconds from the onset of Cue 1 presentation. Classifiers were trained and evaluated at the single-subject level using a four-fold cross validation procedure in which data were randomly partitioned into four subsets. To avoid temporal confounds, all volumes within a trial were kept in the same fold. The accuracies reported are the average of the four folds. For each subject, we tested three classifiers that differed in the spatial filtering of the images used as inputs. The first model used whole-brain images as inputs; feature selection was then performed by selecting 1,000 voxels that were most strongly associated with the training labels based on their ranking of ANOVA F-values. In the second and third models, inputs were constrained to voxels within the aVS and dmVS masks, respectively. To test the significance of classifier performance, we ran permutation tests in which the same classification procedure was repeated using 1,000 random permutations of training labels^57^. This provided a distribution of chance-level accuracy; we consider the performance of a classifier as significantly above chance if is greater than 95% of accuracies obtained using permutated tests^57^.

Additionally, to examine whether individual differences in striatal responses during maze-navigation predicted ventral striatal signals in the effort-based decision-making task, we used the extracted beta parameters from the dmVS (obtained from Navigation Start phase, High-Effort>Low-Effort) and VS (obtained from Reward phase, Reward>No-Reward, High-Effort condition) to compute individual differences in the effect of effort (High-Effort – Low-Effort) during effort initiation for dmVS and during reward receipt for VS. We then correlated these effects with extracted beta parameters from the VS during an open contrast of the presentation of effort information at Cue 1 (averaged across all presented effort levels) in the effort-based decision-making task. An open contrast (i.e., contrasted against the implicit baseline) was used as we felt this best represented neural responses to potential effort as compared to the default “no effort” option that was always available to participants on this task.

