## Supplemental Information for "Distinct striatal subregions and corticostriatal connectivity for effort, action and reward"

**This PDF file includes:**

Tables S1 to S2
Figures S1 to S3

| Regions of Activation | Hemisphere | Peak MNI Coordinates | | | | | |
|---|---|---|---|---|---|---|---|
| | | x | y | z | k | max | mean |
| *Cue phase, High-Effort>Low-Effort* | | | | | | | |
| Precentral Gyrus | Left | -33 | -30 | 60 | 5662 | 8.35 | 4.60 |
| dorsomedial VS | Left | -12 | 0 | 15 | 58 | 6.15 | 4.56 |
| dorsomedial VS | Right | 12 | 12 | 6 | 44 | 4.79 | 4.29 |
| Fusiform Gyrus | Left | -45 | -39 | -9 | 89 | 6.53 | 4.47 |
| Anterior Lobe | Right | 24 | -42 | -30 | 389 | 6.51 | 4.69 |
| Middle Temporal Gyrus | Right | 51 | 9 | -27 | 141 | 6.30 | 4.37 |
| Middle Occipital Gyrus | Right | 36 | -90 | 9 | 341 | 6.26 | 4.48 |
| Cuneus | Left | -21 | -93 | 3 | 419 | 5.88 | 4.52 |
| Midbrain Red Nucleus | Left | -9 | -18 | -3 | 79 | 5.79 | 4.51 |
| Superior Temporal Gyrus | Left | -33 | -54 | 18 | 90 | 5.70 | 4.45 |
| Inferior Semi-Lunar Lobule | Right | 12 | -66 | -48 | 119 | 5.56 | 4.30 |
| Precuneus | Left | -9 | -60 | 30 | 57 | 5.17 | 4.28 |
| *Cue phase, Low-Effort>Passive-Motion* | | | | | | | |
| Postcentral Gyrus | Left | -36 | -24 | 51 | 444 | 6.97 | 4.67 |
| Middle Occipital Gyrus | Right | 33 | -90 | -3 | 68 | 6.59 | 4.67 |
| Superior Parietal Lobule | Right | 21 | -63 | 54 | 104 | 6.15 | 4.65 |
| Superior Frontal Gyrus | Left | -9 | 0 | 63 | 149 | 5.66 | 4.45 |
| Inferior Occipital Gyrus | Left | -30 | -93 | -6 | 82 | 5.25 | 4.29 |
| *Navigation-Start phase, High-Effort>Low-Effort* | | | | | | | |
| Lentiform Nucleus/Putamen | Left | -30 | -18 | -3 | 23080 | 9.74 | 4.61 |
| dorsomedial VS/thalamus | Left | -9 | 0 | 6 | 142 | 6.92 | 4.82 |
| dorsomedial VS | Right | 9 | 6 | -3 | 122 | 6.45 | 4.72 |
| VS | Left | -12 | 15 | -9 | 67 | 5.13 | 4.19 |
| *Navigation-Start phase, Low-Effort>Passive-Motion* | | | | | | | |
| Postcentral Gyrus | Left | -39 | -18 | 51 | 3818 | 16.23 | 6.19 |
| Cerebellum | Right | 21 | -51 | -24 | 934 | 11.11 | 5.41 |
| Midbrain Red Nucleus | Left | -6 | -24 | -9 | 2233 | 10.61 | 5.39 |
| lateral striatum | Left | -24 | 0 | -6 | 411 | 9.26 | 5.78 |
| lateral striatum | Right | 24 | 3 | -3 | 359 | 9.02 | 5.58 |
| Cerebellum | Left | -33 | -51 | -30 | 167 | 6.92 | 4.70 |
| Precuneus | Right | 21 | -54 | 57 | 414 | 6.22 | 4.45 |
| Middle Frontal Gyrus | Right | 33 | 39 | 27 | 63 | 5.68 | 4.48 |
| Cerebellar Tonsil | Left | -18 | -36 | -48 | 53 | 5.12 | 4.08 |
| Middle Temporal Gyrus | Right | 39 | -75 | 9 | 88 | 4.88 | 4.01 |
| Cuneus | Left | -15 | -93 | 6 | 68 | 4.27 | 3.91 |
| *Reward phase, reward>no-reward, all conditions* | | | | | | | |
| VS | Left | -18 | 24 | 0 | 151 | 5.69 | 4.40 |
| VS | Right | 6 | 15 | -3 | 31 | 5.15 | 4.46 |
| VS | Left | -3 | 15 | -3 | 27 | 4.85 | 4.28 |
| *Reward phase, parametric effect of reward magnitude* | | | | | | | |
| VS | Left | -6 | 18 | -6 | 187 | 5.99 | 4.26 |
| VS | Right | 6 | 15 | -3 | 35 | 5.35 | 4.28 |

**Table S1. Results of whole-brain analyses in the maze-navigation task**. Related to Figure 1. Reported clusters survived $p<0.05$ familywise-error correction with voxelwise $p<0.0005$. Local maxima (indented regions) are reported only for striatal sub-clusters. Max and mean t-statistics for each peak and cluster, respectively, are reported. *MNI: Montreal Neurological Institute; VS: ventral striatum.*

| | | Peak MNI Coordinates | | | | | |
|---|---|---|---|---|---|---|---|
| **Regions of Activation** | **Hemisphere** | **x** | **y** | **z** | **k** | **max** | **mean** |
| *Cue 2 > Cue 1* | | | | | | | |
| Lingual Gyrus / Cerebellum | Right | 15 | -81 | -9 | 318 | 7.62 | 5.04 |
| Postcentral Gyrus | Left | -57 | -21 | 21 | 516 | 6.91 | 4.69 |
| Middle Occipital Gyrus | Right | 51 | -75 | 6 | 111 | 6.78 | 4.84 |
| Precentral Gyrus | Left | -45 | 0 | 6 | 311 | 6.59 | 4.51 |
| Lingual Gyrus / Cerebellum | Left | -27 | -69 | -21 | 143 | 6.17 | 4.58 |
| dACC | Left | -6 | -3 | 48 | 160 | 5.77 | 4.34 |
| *Cue 2, parametric effect of SVC* | | | | | | | |
| dACC | Left | 0 | 18 | 54 | 122 | -4.82 | -4.24 |

**Table S2. Results of whole-brain analyses in the effort-based decision-making task**. Related to Figure 3. Reported clusters survived $p<0.05$ familywise-error correction with voxelwise $p<0.0005$. Max and mean t-statistics for each peak and cluster, respectively, are reported. *MNI: Montreal Neurological Institute; SVC: subjective value of chosen option; dACC: dorsal anterior cingulate cortex*.

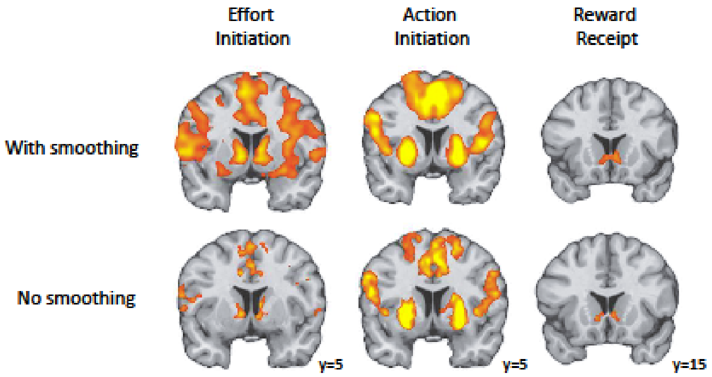


**Figure S1. Activation of striatal subregions in smoothed and unsmoothed data**. Related to Figure 1. All maps are familywise-error corrected at $p<0.05$ using voxelwise threshold of $p<0.0005$. The effort initiation (High-Effort>Low-Effort) and action initiation (Low-Effort>Passive-Motion) contrasts are examined during the initiation of navigation (Navigation Start phase), and the reward receipt contrast (reward>no-reward, across all conditions) is examined during the Reward phase of the maze-navigation task.

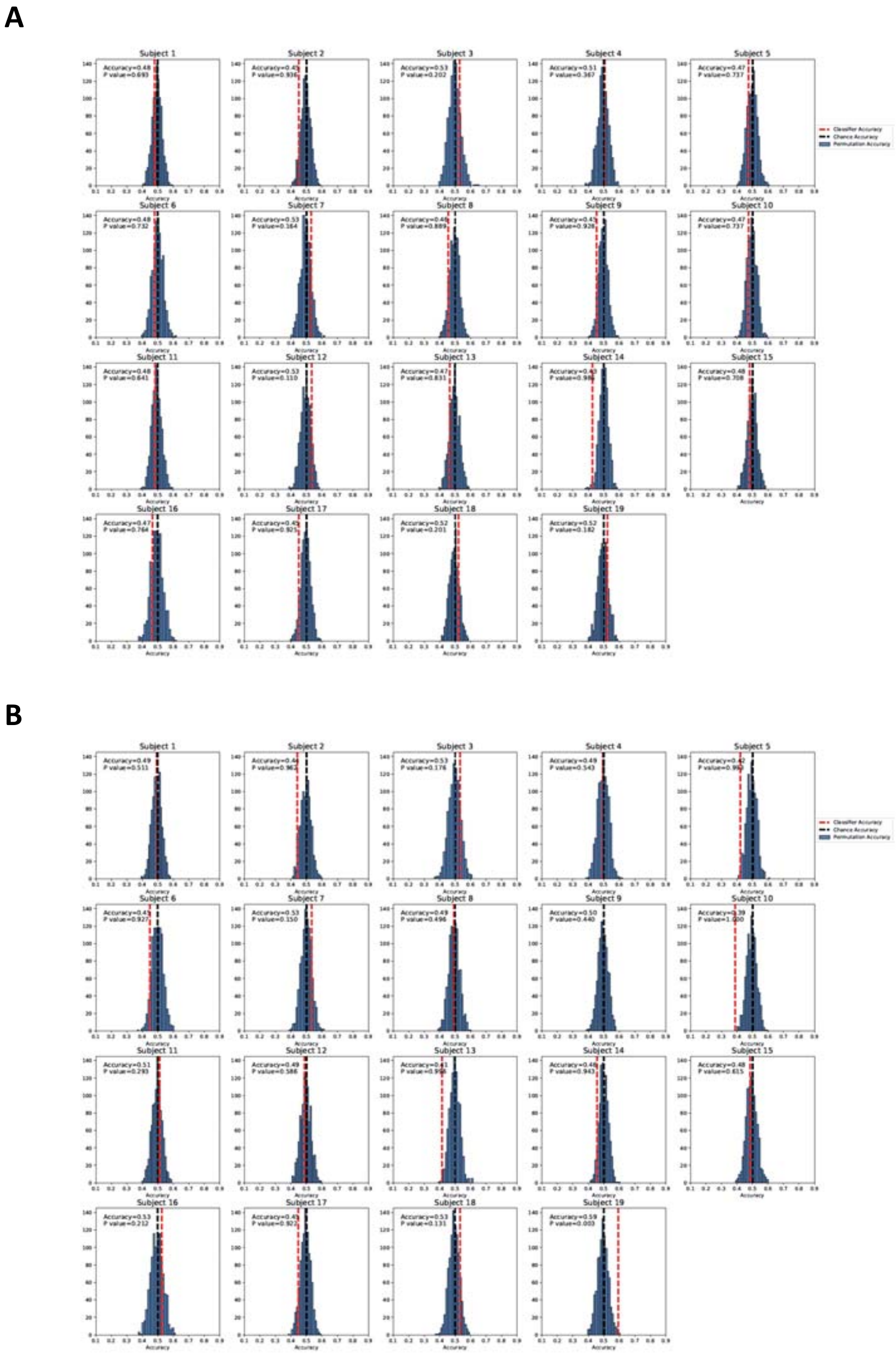

**Figure S2. MVPA results using voxels within striatal subregions to predict the value information presented during the effort-based decision-making task.** Related to Figure 3. Plots show distributions of accuracies from 1000 permutation tests for each participant for classifiers trained on voxels within (A) dmVS (group-level accuracy: M=0.48, std=0.05) and (B) aVS (group-level accuracy: M=0.48, std=0.03).

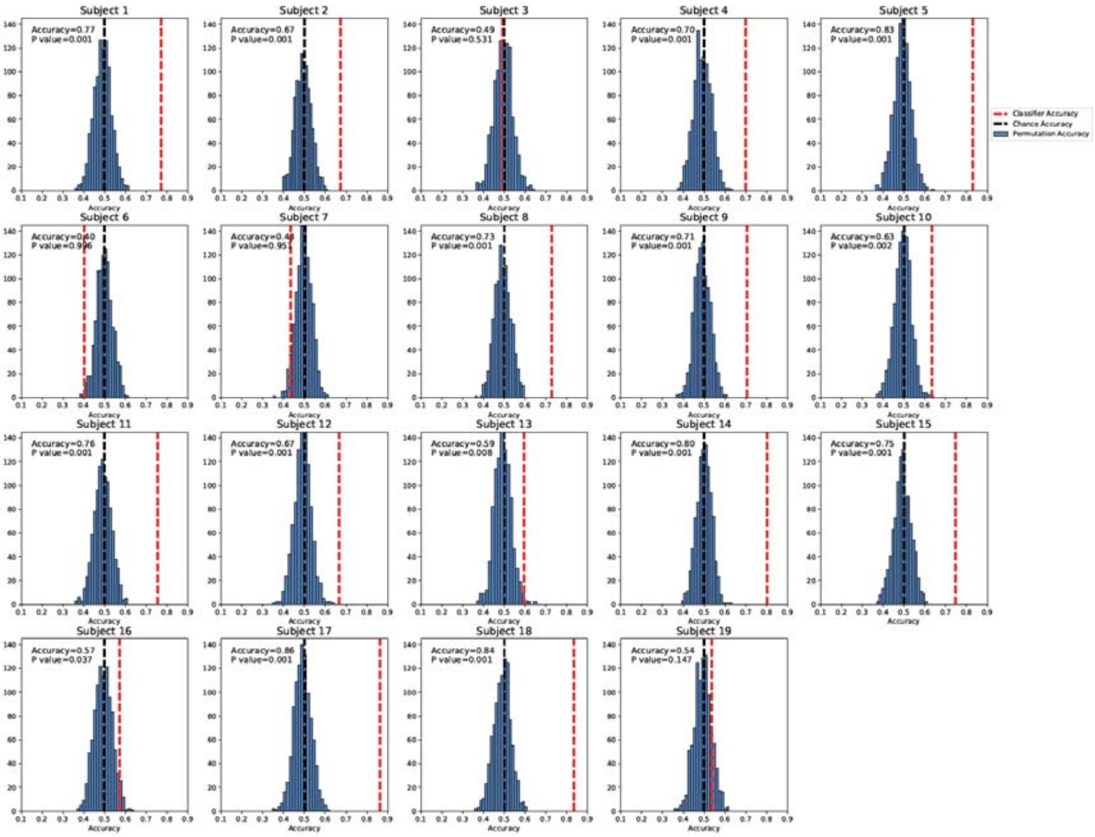


**Figure S3. MVPA results using all voxels within the brain to predict the value information presented during the effort-based decision-making task**. Related to Figure 3. Distributions of accuracies from 1000 permutation tests for each participant. This model predicted the type of value information above chance level for 78.9% of individuals (group-level accuracy: M=0.67, std=0.14).
